# Much more than a clasp: Evolutionary pattern of amplexus diversity in anurans

**DOI:** 10.1101/854117

**Authors:** Juan D. Carvajal-Castro, Yelenny López-Aguirre, Ana María Ospina-L, Juan C. Santos, Bibiana Rojas, Fernando Vargas-Salinas

**Author notes:** **Corresponding author:** Juan D. Carvajal-Castro, postal address: Cll 22c # 29a-47, Bogotá, Colombia.

## Abstract

The evolution and diversification of animal reproductive modes have been pivotal questions in behavioral ecology. Amphibians present the highest diversity of reproductive modes among vertebrates, involving various behavioral, physiological and morphological traits. One of such features is the amplexus, the clasp or embrace of males on females during reproduction, which is almost universal to anurans. Hypotheses about the origin amplexus are limited and have not been thoroughly tested, nor had they taken into account evolutionary relationships in most comparative studies. However, these considerations are crucial to understand the evolution of reproductive modes. Here, using an evolutionary framework, we reconstruct the ancestral state of amplexus in 686 anuran species; investigate whether the amplexus type is a conserved trait; and test whether sexual size dimorphism (SSD) could have influenced the amplexus type or male performance while clasping females. Overall, we found evidence of at least 35 evolutionary transitions in amplexus type across anurans. We also found that amplexus exhibits a high phylogenetic signal (it is conserved across Anura evolutionary history) and the amplexus type does not evolve in association with SSD. We discuss the implications of our findings on the diversity of amplexus types across anurans.

## INTRODUCTION

Understanding the evolution and diversification of reproductive modes in animals has been a shared interest among evolutionary biologists over decades (e.g., Salthe, 1969; Shine, 1983; Craig, 1987; Alves *et al*., 1998; Blackburn, 2000; Crespi & Semeniuk, 2004; Haddad & Prado, 2005). In addition to sexual selection, natural selection promotes reproductive diversity by favoring modes of reproduction that maximize the likelihood of successful matings in a given environment (Pianka, 1976; Zamudio *et al*., 2016). Thus, reproductive modes are one of the most critical life-history traits directly affecting fitness and survival via physiological, morphological and behavioral adaptations that increase each individual’s ability to find mates and producing viable offspring, in response to environmental (and other) selective pressures (Angelini & Ghiara, 1984; Lodé, 2012).

Anuran amphibians exhibit one of the highest diversities in reproductive modes among vertebrates (Duellman & Trueb, 1986; Vitt & Caldwell, 2014). These reproductive modes are defined as a combination of ecological, developmental and behavioral traits that include oviposition site, ovule morphology, clutch size, developmental rate, and presence or absence of (different types of) parental care (Salthe & Duellman, 1973; Duellman & Trueb, 1986). Thus, anuran reproductive modes exhibit a gradient of parental involvement which ranges from no or little parental care, involving mostly aquatic oviposition of clutches with hundreds or thousands of eggs; to elaborate parental care with relatively few terrestrial eggs, decisive parental involvement or direct development with reduced or absent tadpole stage (Hödl, 1990; Haddad & Prado, 2005; Crump, 2015).

Previous studies have investigated different aspects of the evolution of reproductive modes in anurans using limited phylogenetic comparative methods (Duellman, 2003; Gomez-Mestre *et al*., 2012; Zamudio *et al*., 2016; Furness & Capellini, 2019). These studies have hypothesized that ecological and population structure factors such as desiccation in temporary ponds, availability of humid microhabitats in terrestrial environments and predation are major selective forces shaping most reproductive modes. However, to our knowledge, none of these studies has addressed the evolution of a key behavioral component in frog reproduction: the amplexus or ‘mating clasp’. Here, we investigate the evolutionary patterns of this trait across the Anura and reveal major evolutionary transitions in this crucial component of frogs’ reproductive behavior.

Amplexus is present in most anuran species and consists of a male grasping a female from behind with his forelimbs. Thus, not surprisingly, it has been interpreted as a behavior by which a male ensures the fidelity of its female partner during mating, increasing the chance of egg fertilization (Duellman & Trueb, 1986; Wells, 2007). Like other mating traits in anurans, amplexus types are diverse see Fig. 1). For instance, the inguinal amplexus is considered a basal condition to all anurans, while the axillary amplexus and its variations (including the lack of amplexus altogether) are considered as derived states (Duellman & Trueb, 1986; Wells, 2007; Pough *et al*., 2016). Several hypotheses have been proposed to address the evolution of such amplexus diversity, suggesting that that variants of the amplexus may have evolved as a consequence of sexual size dimorphism, parental care and the ecological factors affecting oviposition site (Duellman & Trueb, 1986; Wells, 2007; Pough *et al*., 2016). However, these ideas have not been tested under a phylogenetic framework. Such comparative analyses could greatly improve our understanding of the evolutionary patterns of anuran amplexus diversity, and offer new baseline data for further comparative studies about the behavioral ecology of reproductive modes among vertebrates.

**Figure 1.**
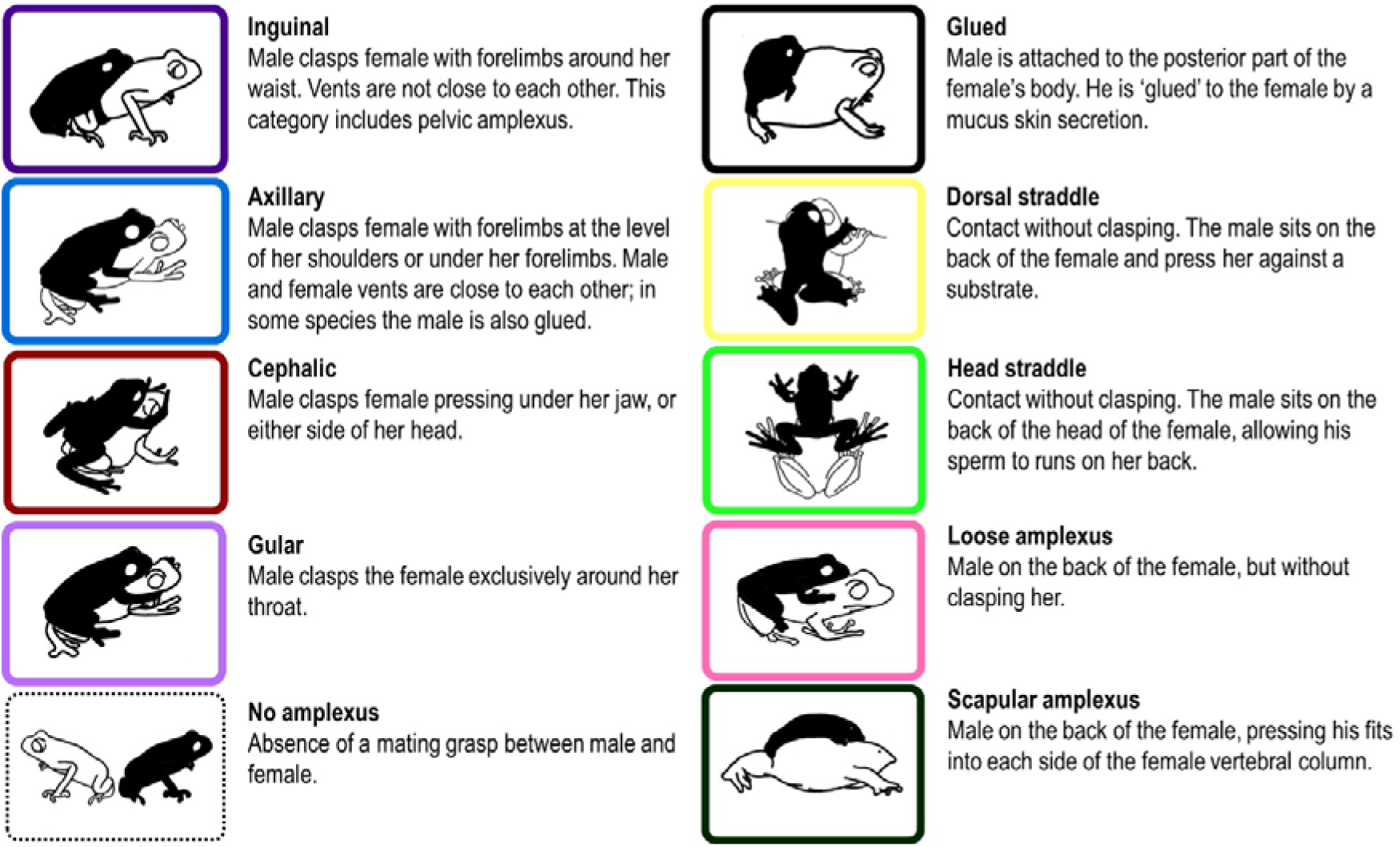
Diversity of amplectic positions in anurans (males in black color). Definitions according to Blommers-Schlosser (1975), Townsend & Stewart (1986), Duellman & Trueb (1986), Wells (2007), Zachariah *et al.,* (2012), Jungfer *et al.,* (2013) and Willeart *et al*., (2016). Because descriptions of some amplexus types are not clear or are ambiguous, we pooled the category “Independent” (cited by Duellman & Trueb, 1986), as “No amplexus”. Frame colors make references to the same amplexus types shown in Fig. 2 and 3.

Here, we use the most complete phylogeny of Anura (Jetz & Pyron, 2018) to map the origins and diversification of the known types of amplexus in 686 species with reported records. Furthermore, we explore the relationship between the evolution of amplexus diversity and sexual dimorphism in body size (measured as female-to-male snout-vent length ratio, hereafter SSD). We predict that species with a male-biased, little or no SSD, would benefit from axillary amplexus, whereas species with a high female-biased SSD would likely present inguinal amplexus or another derived type of amplexus or strategy (e.g. ‘glued’ in Fig. 1). These predictions are based on the physical restrictions that a very small male could have to clasp a large female and, hence, to ensure her fidelity during mating. Overall, our results show that the different types of amplexus, and the lack thereof, are well-defined throughout the Anura tree of life. This is a fundamental step forward to understanding how environmental factors and life history have shaped the amazing diversity of reproductive modes across Anura and other vertebrates.

## MATERIAL AND METHODS

### AMPLEXUS CHARACTERIZATION

We looked for information on types of amplexus in Anura from primary literature (e.g., peer-reviewed articles, books) located by using Google Scholar and Web of Science (WoS). We used “amplexus” and “nupcial clasp” as keywords. Because we obtained an excess of results unrelated to Anura (the term amplexus has also been used for invertebrates; Conlan, 1991), we included the keywords “anura” and “frogs”. To further narrow down our search, we also combined previous keywords with anuran families’ names (e.g., “Amplexus” AND “Dendrobatidae”). Within the selected publications, we searched for an account describing male and female behavior with enough detail (e.g. an observational account of the behavior or photograph) to be assigned to an amplexus type.

We defined the documented amplexus types (Fig. 1) following Duellman & Trueb (1986) and Willaert *et al*., (2016), but also considering the following clarifications: First, in several species it has been reported that the amplexus type might change at the moment of oviposition (e.g. *Anomaloglossus bebei* and *Brachycephalus ephippium*; Bourne *et al.,* 2001, Pombal *et al*., 1994; see supplementary material for more examples). In these cases, we considered the preoviposition amplexus type to be the predominant one (i.e. longer duration and most frequently reported in literature, as in most cases researchers did not wait until oviposition for recording breeding behavior). Second, some studies include observations of multiple types of amplexus for a given species. In these cases, we used the report(s) with the strongest evidence, which include a textual description of the type of amplexus or visual evidence such as photographs or videos. Below, we provide a list of some specific examples of conflicting reports and evidence of new types of amplexus.

For *Nyctibratrachus humayumi* (Nyctibatrachidae), it was reported that this species lacks amplexus (Kunte, 2004), but it was later observed that *N. humayumi* has a dorsal straddle amplexus (Willaert *et al.,* 2016). In another case, the authors cite a pectoral amplexus for *Nasikabatrachus sahyadrensis* (Sooglossidae) (Zachariah *et al.,* 2012), but based on the detailed description by the authors, we consider this behavior to be a new type of amplexus. We define this clasp as a ‘scapular amplexus’, where the male presses his fists into each side of the female’s vertebral column. For *Scaphiophryne gottebei* (Microhylidae), the authors mention an inguinal amplexus, but this account includes several figures showing a characteristic axillary amplexus (Rosa *et al.,* 2011). For *Osteocephalus* (Hylidae), we used the ‘gular amplexus’ definition by Jungfer *et al.,* (2013), which refers to a type of amplexus where the male clasps the female exclusively around her throat. Likewise, some species with axillary amplexus have been reported to have a ‘glued amplexus’ (e.g. *Elachistocleis bicolor* (Cacciali, 2010) and *Chiasmocleis leucosticta* (Haddad & Hödl, 1997)). Such reports indicate that the males are ‘glued’ to the female dorsum but, in our analysis, we did not take this clasping behavior as different from an axillary amplexus. Our consideration is based on the fact that males of species with axillary amplexus have not been examined thoroughly enough for the presence or absence of glands to suggest that these organs exist, or produce sticky or glue-like substances. For species like *Nyctibatrachus aliciae* (Nyctibatrachidae, Biju *et al.,* 2011) or *Mantella aurantiaca* (Mantellidae, Vences, 1999, Glaw & Vences, 2007), the observed amplexus consists of a male sitting on the dorsum of the female by a short period of time, without an actual clasp; we classified such observations as ‘loose amplexus’. Lastly, *Aplastodiscus leucopygius* (Hylidae) and *Ascaphus truei* (Ascaphidae) have a ‘dynamic amplexus’, which is difficult to categorize because at different moments the same pair (i.e. male and female) exhibits diverse amplexus positions (Stephenson & Verrell 2003; Berneck *et al*. 2017). Therefore, these two species were not included in our analysis.

We completed the life history characterization for all species with amplexus data by including male and female body size (i.e. snout-vent-length, SVL) and sexual size dimorphism (SSD). In cases where only the range of body size was available, we used the median values.

### COMPARATIVE METHODS

For the phylogenetic analysis, we obtained 1000 random phylogenies based only on genetic data from Jetz & Pyron (2018). Then, we ran the analysis 1000 times to evaluate the robustness and include uncertainty in topologies in the R software environment (R Core Team, 2018). Later, we made a stochastic ancestral reconstruction (Bollback, 2006) of the character “type of amplexus” using 1000 trees, with the *make.simmap* function in ’phytools’ R-package version 0.6-99 (Revell, 2012) which is a best-fit character evolution model (Only with 100 trees). We contemplate an “Equal Rate model” (ER), which assumes that all transitions between traits occur at the same rate (Pagel 1994; Lewis 2001); and an “All-Rate-Different model” (ARD), which assumes that all transitions between traits occur at different rates (Paradis *et al*., 2004). We used the *fitDiscrete* function in the ’geiger’ R-package (v. 2.0.6.2; Harmon *et al.,* 2007) to compare the ER and ARD models, and selected the model with the lowest AICc value. We did not generate fully-sampled phylogenies using the taxonomic imputation method to make our ancestral character reconstruction because this approximation has been demonstrated to be inappropriate for this kind of analysis due to increased bias (Rabosky, 2015; Rocha *et al.,* 2016; Jetz & Pyron 2018).

To test whether the type of amplexus and SSD are conserved or labile (convergent) traits, we calculated their phylogenetic signal. This property is defined as the tendency of traits in related species to resemble each other more as a consequence of shared ancestry (Blomberg & Garland 2003). For this purpose, we used the statistic lambda (λ) proposed by Pagel (1997, 1999) as a measurement of phylogenetic signal. The λ value varies from 0 to 1; if λ ∼ 1, it indicates a strong phylogenetic signal (i.e. conserved trait), whereas if λ ∼ 0, it indicates that the evolution pattern of the trait has been random or convergent; that is, these characters lack phylogenetic signal (Gómez *et al.,* 2010, Kraft *et al.,* 2007; Revell *et al.,* 2008). Calculation of the phylogenetic signal for type of amplexus was made using the *fitDiscrete* function of ’geiger’ R-package. In addition, we calculated the likelihood of a model with no phylogenetic signal and the maximum likelihood value ofλ; later we used a likelihood ratio test to compare these two models and calculate a p-value (significance alpha 0.05) under a chi-square distribution. For SDD, phylogenetic signal was tested with the *phylosig* function in ’phytools’ (Revell, 2012).

To test whether the evolution rates of type of amplexus have increased or slowed over time, we used a delta model in *fitDiscrete* function of ’geiger’. In addition, we calculate the likelihood of a model with delta equal to one and a model with observed delta; later we used a likelihood ratio test to compare these two models and calculated a p-value (significance alpha 0.05) under a chi-square distribution. If delta statistic > 1, this indicates that recent evolution has been relatively fast; in contrast, if delta ≤ 1, it indicates that recent evolution has been relatively slow. Because the difference in body size between males and females could promote changes in types of amplexus due to mechanic incompatibility (e.g., small males might not physically clasp a large female), we used a phylogenetic ANOVA to compare sexual size dimorphism across different types of amplexus. This analysis was performed using the *phylANOVA* function in ’phytools’.

## RESULTS

Our analyses included 686 species from 46 anuran families (Table S1). The distribution of species in our dataset comprised all continents where amphibians are present. Most families (i.e. 35 families) have only one type of amplexus, while 11 families have more than one (Fig. 2, Table S1). The average SSD is 1.17 ± 0.16 (range = 0.70-1.85; n= 478 species), and it varies between anuran species and families (Fig. 2, Table S1); in 42 species (8.78%) male body size is larger than female body size, while in most cases (429 species, 89.74%) the female is larger than the male; in the remaining seven species (1.46%) males and females exhibit similar body size.

**Figure 2.**
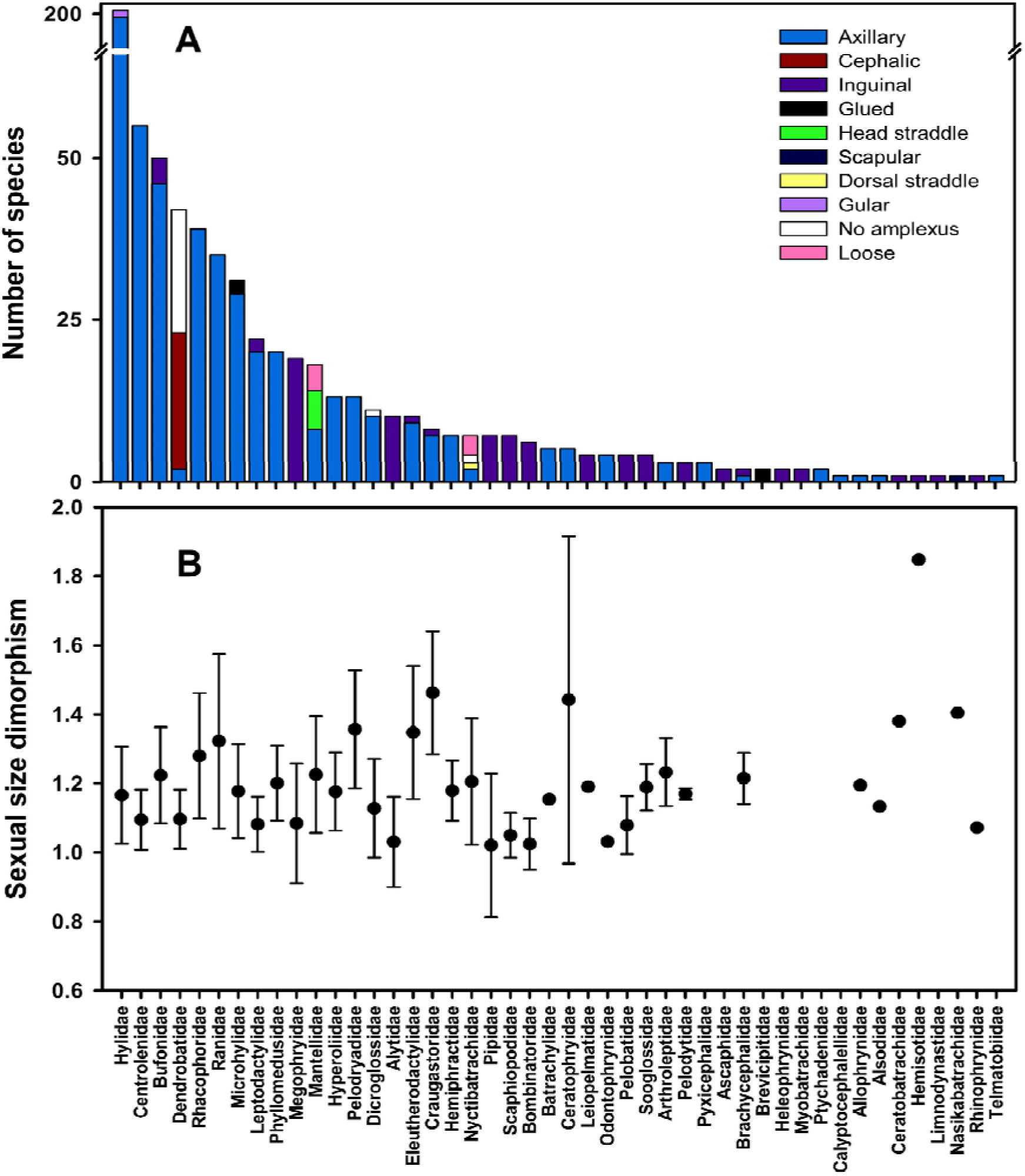
**A**. Summary of the number (686) of anuran species, type of amplexus (or lack thereof), and **B.** sexual size dimorphism (female-to-male snout-vent length ratio; SSD) per Family of Anura included in this study. Plot based on data from Table S1. Dots and bars in the plot B indicate mean values and standard deviation, respectively; for some families was not possible to calculate SSD because absence of data for female or male body size.

For the ancestral reconstruction of types of amplexus, we chose the ER model (*LnL*= −227.438, *AIC*=456.882) over ARD model (*LnL*=-165.484, *AIC*= 538.497) based on the lowest AICc value (Fig. 3). Our results support the inguinal amplexus as the basal state to all Anura. For instance, Ascaphidae and other basal frog families (e.g., Leiopelmatidae, Bombinatoridae, Alytidae, Pipidae) present inguinal amplexus. The axillary amplexus was found to be the most frequent state, occurring in 540 out of the 686 species (i.e. 78.72%). However, we found 35 evolutionary transitions between all type of amplexus across the whole Anura phylogeny (Fig. 4A). The greatest number of evolutionary transitions (i.e., 12) occurred between axillary and inguinal amplexus states (Fig. 4A). Likewise, we found a strong phylogenetic signal amplexus (Page’s λ = 0.963, *P* < 0.0001). In contrast, we found that SSD has a weak phylogenetic signal (Page’s λ = 0.266, *P* = 0.043). We also found that the delta value is equal to 0.694 (SD = 0.208), but when we compare it with a model with delta equal to 1, we found no differences (*P* = 0.63). Furthermore, we did not find differences in SSD across types of amplexus (*F* = 1.2086, *P* = 0.4215, n= 478; Fig. 4B).

**Figure 3.**
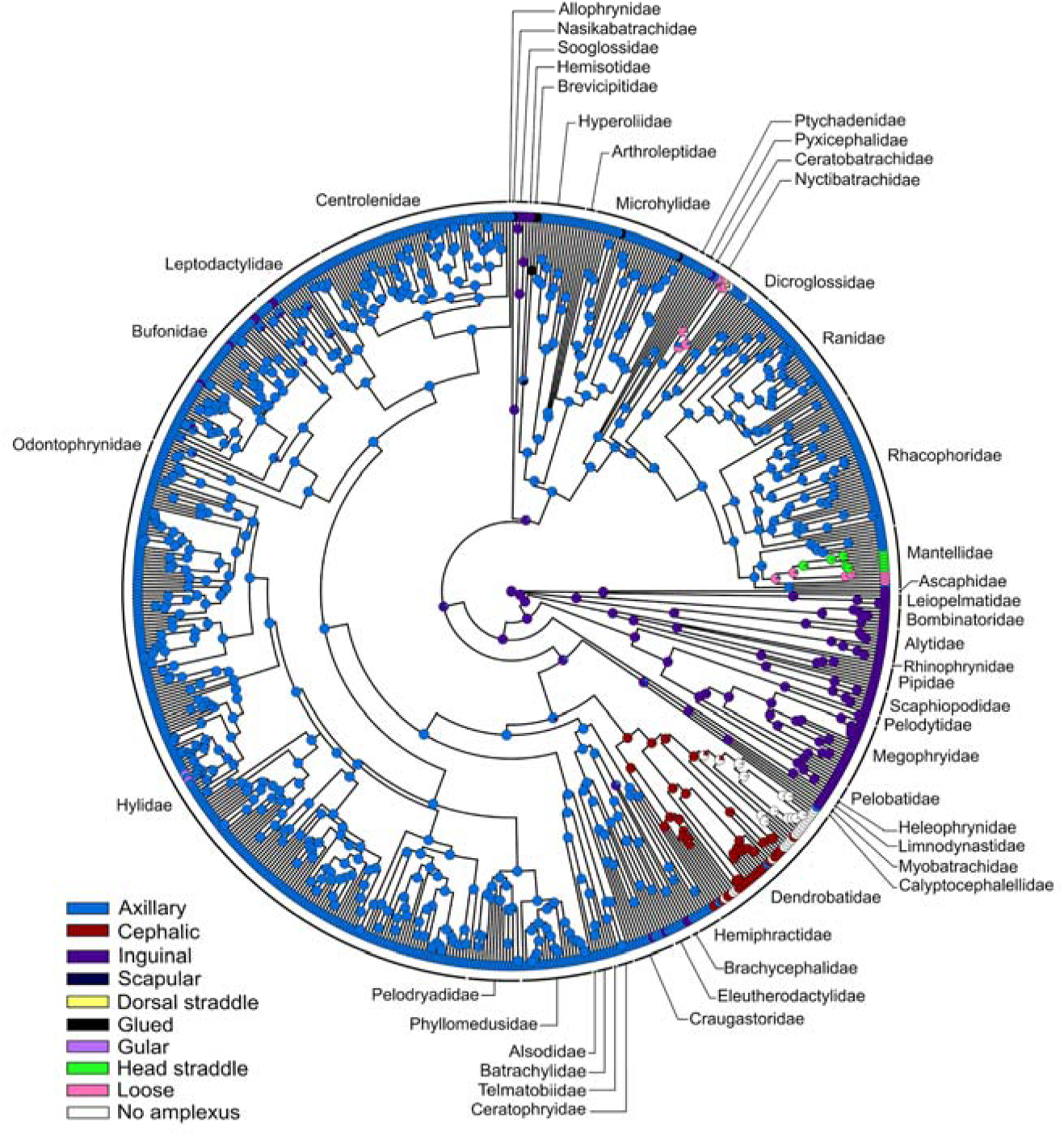
Ancestral reconstruction of amplexus type using 1000 trees for 686 anuran species under an “equal rates” model of trait evolution. Tips represent actual type of amplexus for each species and nodes represent the probability of each type of amplexus (Table S2 for specific values). Phylogenetic tree from Jetz & Pyron (2018).

**Figure 4.**
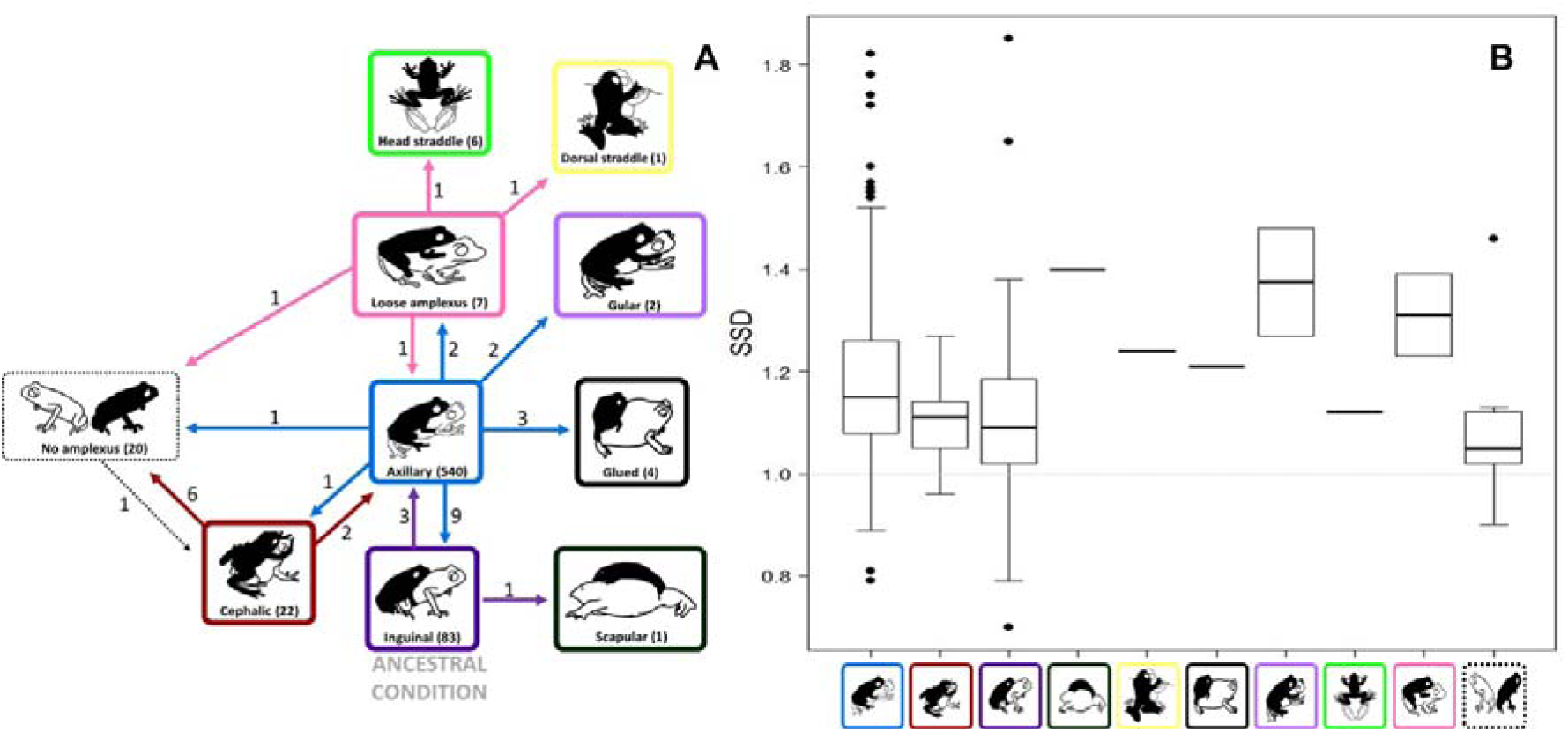
**A.** Estimated number of independent evolutionary transitions in type of amplexus for the 686 anuran species included in this study. In parentheses are pointed the number of species with each type of amplexus. Estimates are based on 1000 phylogenetic trees, and an ancestral state reconstruction performed with phytools packages (see Fig. 2 and text for details). **B.** Variation in sexual size dimorphism (female-to-male snout-vent length ratio, SSD) in 478 species with different types of amplexus. Phylogenetic ANOVA: (*F* = 1.2086, *P* = 0.4215, n= 478).

## DISCUSSION

Different selective pressures are known to shape the behavioral, physiological and physical traits that characterize the diverse reproductive modes and behaviors in anurans (and fishes) in comparison to other vertebrates. One of s traits is the amplexus, whose evolutionary trends we characterized using a comparative phylogenetic framework. For this purpose, we explored the relationship between the different types of amplexus (or lack thereof) and sexual size dimorphism. Below, we discuss the implications of our findings and generate testable hypotheses for future research on the evolution of reproductive modes in anurans.

We found a significant phylogenetic signal in amplexus. This result was anticipated, considering that the amplexus is an aspect of the anuran reproductive behavior that is expected to have been shaped by other various factors not related to reproduction (e.g. morphology, microhabitat; Duellman and Trueb 1986, Wells 2007). Likewise, it is presumed that the type of amplexus affects the reproductive success of males and females in several ways. For example, the type of amplexus determines the proximity of cloacas between males and females and, possibly, the success of egg fertilization (Davis and Halliday 1979; Wells 2007). Altogether, we infer that any change between types of amplexus requires several selective factors to act in tandem and weighed by their combined selective force. While we found a slight indication of a slow down in the rate evolution of amplexus (delta value equal to 0.694), this was not supported when we tested if this parameter estimate was different from delta = 1. In other words, the amplexus rate of change follows a Brownian motion model which does not support a slow-down, but rather a random change (either slow-down or acceleration) in its rate of evolution through the evolutionary history of Anura.

In contrast to amplexus type, the weak phylogenetic signal in SSD suggests that selective forces might be promoting body size disparities to be the result of convergent evolution across Anura. For instance, in most frogs, females are larger than males because there is a strong positive relationship between fecundity and body size (i.e., selection favors larger size in females); in contrast, a larger body size in males is not necessarily tied to higher attractiveness to females or higher dominance across species (Halliday and Tejedo 1995; Wells 2007).

Previous reports support our results that all basal lineages (e.g., Ascaphidae, Pipidae, and Myobatrachidae) have an inguinal amplexus (Lynch, 1973; Rabb, 1973; Weygoldt, 1976), which we found to be the ancestral state across anurans. Likewise, axillary amplexus appears to have been derived from the inguinal type, as hypothesized by others (Duellman & Trueb, 1986; Wells, 2007; Vitt & Caldwell, 2014; Pough *et al*., 2016). Most importantly, our analysis revealed that at least 35 transitions have occurred between amplexus types across the anuran phylogeny. Notably, we found that transitions from axillary to other type(s) of amplexus have occurred in high frequency (i.e., at least 18 times). These results suggest that axillary amplexus might be a key intermediate type that eventually diversified in almost all other amplexus types.

Axillary amplexus is the most widespread type across Anura and, according to our analysis, it evolved at least three independent times from the ancestral inguinal amplexus. The prevalence of axillary amplexus suggests its high versatility in different ecological contexts that can effectively relate to the reproductive success of males. For instance, in many species, especially those denominated by explosive breeders where males and females congregate to reproduce (Wells, 1977), it is common to find uncoupled males trying to get off amplectant males from the dorsum of their partners (Halliday & Tejedo, 1995; Wells, 2007; see video S1 for an example in *Rhinella castaneotica* (Caldwell, 1991)). In this context, amplectant males would benefit from grasping the female as strongly as possible and only the axillary amplexus is the most effective (see Lee & Corrales, 2002; Vargas-Salinas, 2005 for an example with *Rhinella marina* (Linnaeus, 1758)). Intrasexual competition between males may also favor axillary amplexus where the intense competition during short breeding seasons results in the evolution or persistence of traits that reinforce mate guarding behavior by males (e.g. axillary amplexus plus been glued). In contrast, most other types of amplexus might be related to taxa with prolonged breeding seasons where amplectant males are exposed to lower risks of being displaced by competing males (Duellman & Trueb, 1986; Wells, 2007; Willeart *et al*., 2016).

If the evolution of the axillary amplexus has been so successful, and this type of amplexus is so versatile, what could promote the evolution of other types of amplexus? Furthermore, why do reversions to the inguinal amplexus occur? Surely, the type of amplexus in a given species or clade has been shaped by multiple environmental, physiological and morphological factors and particular social contexts (see possible scenarios below). This pattern is also true for other aspects of reproductive modes, and likely applicable to vertebrate groups other than amphibians. Testing all the possible scenarios is beyond the scope of this study, but we propose some inferences, as follows. For example, effective antipredator chemical defenses (i.e., toxic alkaloids) such as those in Dendrobatidae (poison frogs) can promote amplexus diversification. Several lineages of poison frogs have evolved aposematic coloration (Santos *et al*., 2003; Rojas, 2017), which is associated with a high diversification in acoustic communication signals as an alleged indirect effect of a reduction in predation pressure (Santos *et al*., 2014); thus, aposematism could also allow an increase in the complexity of courtship behaviors, promoting matings where axillary amplexus is not necessary. Our results support such intuition, as at least 22 dendrobatid species exhibit cephalic amplexus, whereas 18 species exhibit no amplexus (Weygoldt, 1987; Castillo-Trenn & Coloma, 2008). Moreover, most species of Dendrobatidae are prolonged breeders (Wells, 1977), mostly terrestrial, highly territorial and whose oviposition occurs in hidden places under leaflitter and tree roots (Wells, 1978; Pröhl, 2005; Summers & Tumulty, 2014; Rojas & Pašukonis, 2019). Under these conditions, it might be assumed that aposematic males have fewer risks of predation and losing a female, attracted via acoustic and visual signals, because of the action of an intruder male (Zamudio *et al*., 2016). Thus, aposematic species could evolve complex mating behaviors and diversity of amplexus types if the cost of predation is minimized. In contrast, species that rely on avoiding detection by predators may have evolved mating strategies that offer a balance between attracting females and avoiding enemies.

Microhabitat, or the environment context where males court, is an important factor often overlooked in discussions about the evolution of amplexus diversity. Axillary amplexus may function well in diverse microhabitats (arboreal, terrestrial, aquatic; see Fig. S1). In arboreal species, for example, when a female jumps from leaf to leaf or across branches, a male clasping her in an axillary amplexus would not prevent her from achieving the highest jumping performance; thus this kind of amplexus would be selectively advantageous. A similar performance in arboreal microhabitats would be difficult for inguinal or cephalic amplexus, as the movement of the male’s body during jumping would be erratic and unbalanced with respect to that of the female’s. This hypothesis warrants further investigation from the perspective of the functional association, e.g., between locomotor performance and a specific amplexus type in a particular microhabitat. Likewise, performance experiments would be required to account for the diameter, inclination and type of substrate, which significantly affect the kinematics of locomotion and thus select for specific morphologies and behaviors across a variety of taxa (Andersson, 1994; Losos, 2009; Herrel *et al.,* 2013).

We found a few instances of reversal from the axillary to the inguinal amplexus. A possible explanation for such transitions is that inguinal amplexus promoted morphological adaptations related to thermoregulation (Ashton, 2002; Meiri & Dayan, 2003; Zamora-Camacho *et al.,* 2014). Alternatively, the evolutionary interpretation of such rare transition may be an adaptation to fossorial habits (e.g., Microhylidae and Hyperoliidae). In animals with inguinal amplexus, this behavior allows the female to avoid digging a wider burrow on the ground (Duellman & Trueb, 1986). We hypothesize that this mechanical limitation might explain the evolution of this type of amplexus in *Osornophryne*, a genus of toads with relatively short limbs and distributed in middle and high elevations in the Andes of South America (Frost, 2019). As these species are adapted to cold climates, their globular body shape and short legs require a type of amplexus that provides a better grasping potential for males.

Contrary to our predictions, we found no relationship between sexual size dimorphism (SSD) and type of amplexus. Differences in body size between the sexes can impose physical restrictions to males for clasping females in a way that cloacas are aligned and egg fertilization is optimized (Davis & Halliday, 1977; Ryan, 1985; Robertson, 1990; Bourne, 1992). Moreover, differences in body size between sexes could reduce the strength with which a male can clasp a female, hence reducing a male’s likelihood of being displaced by competing males (Brunning *et al*., 2010). It is possible that amplexus type is related to sexual dimorphism in body shape, or the interaction between body shape and size, rather than size dimorphism alone. For instance, in species with globular bodies and short limbs (e.g. genus *Breviceps*; *Nasikabatrachus sahyadrensis*), the axillary amplexus is mechanically less feasible. In such species, the male is often very small with respect to the female and, thus, the evolution of “alternative strategies” to enhance amplexus may have been advantageous. We propose that the evolution of mucus skin secretions could have increased the effectiveness of gamete transfer in the absence of an actual clasp.

Compared to other amphibian orders, our results reveal some interesting differences. For instance, a recent study involving 114 salamandrid species reports that the ancestral states for this clade were mating on land, oviparity and absence of amplexus (Kieren *et al*., 2018). The authors further suggest that the presence or absence of amplexus might be unrelated with the mating habitat. Our results suggest that anurans exhibit many more reproductive modes as a consequence of a higher species diversity, diverse morphology and more diverse reproductive strategies (Hutter *et al*., 2017; Vitt & Caldwell, 2014; Frost, 2019). Furthermore, this diversity increases towards tropical regions (e.g., Duellman, 1988; Hödl, 1990; Haddad & Prado, 2005). Therefore, complex habitats such as the tropics may offer more opportunities for adaptation in the context of reproductive characteristics, where environmental conditions affect the evolutionary patterns of amplexus diversity. This habitat complexity may have also influenced the reproductive mode diversity found in other ectothermic vertebrates.

The type of amplexus in anurans, or the lack thereof, is related to behavioral features that clearly can affect the reproductive success of individuals (Duellman & Trueb, 1986, Wells, 2007; Buzatto *et al*., 2017). Our findings highlight not only the value of implementing phylogenetic comparative approaches for recording the evolutionary history of reproductive traits in vertebrates, but also the importance of doing detailed field observations of reproductive behavior and natural history. Precisely the lack or infrequency of such kind of reports is one of the main difficulties faced by researchers aiming to do analysis like ours on amphibians and other taxa. Surely, the diversity of amplexus and associated behaviors is much higher than what has been reported to date in the scientific literature, not only among species but even at intraspecific level (Pombal *et al.,* 1994; Stephenson & Verrell, 2003; Berneck *et al.,* 2017). Our study represents a unique large dataset on amplexus types for anurans, and allows to highlight three amplexus types (loose amplexus, gular amplexus, scapular amplexus) that have been overlooked in key literature references (i.e. Duellman & Trueb, 1986; Wells, 2007; Vitt & Caldwell, 2014; Pough *et al*., 2016; Willaert *et al.,* 2016). We hope that further studies about breeding behavior in anurans include detailed observations and descriptions that could reveal novelty aspects associated to diversity of breeding strategies in vertebrates, even in those lineages considered as well studied.

## ACKNOWLEDGEMENTS

JDCC and AMOL thank the Humboldt Institute for logistical support, FVS and YLA are grateful to Universidad del Quindío for support, BR is funded by the Academy of Finland (Academy Research Fellowship No. 21000042021), JCS was supported by start-up funds from SJU. Special thanks to M.P. Toro for help with Figure 2. We are grateful to J. Valkonen, S. Casas, L.A. Rueda-Solano, J. A. Allen and two anonymous reviewers for general feedback and constructive comments on previous versions of this manuscript.

## Supplementary data

Table S1 and references, Video S1, and table S2 can be found in DOI: 10.6084/m9.figshare.11050685

---

**Figure S1.** Axillary amplexus in species under different microhabitat conditions (Arboreal, Terrestrial, Aquatic). **A.** *Atelopus favescens* (Bufonidae), B. *Centrolene savagei* (Centrolenidae), **C.** *Ceratophrys calcarata* (Ceratophrydae), **D.** *Dendropsophus triangulum* (Hylidae), **E.** *Engystomops pustulosus* (Leptodactylidae), **F.** *Agalychnys callidryas* (Phyllomedusidae), **G.** *Pristimantis orpacobates* (Pristimantidae), **H.** *Lithobates vaillanti* (Ranidae). Pictures by B Rojas (A), F Vargas-Salinas (B,D,E,G,H), LA Rueda-Solano (C), and AM Ospina-L (F).

