## Supplementary figures and images for "Much more than a clasp: Evolutionary pattern of amplexus diversity in anurans"

### Figure S1

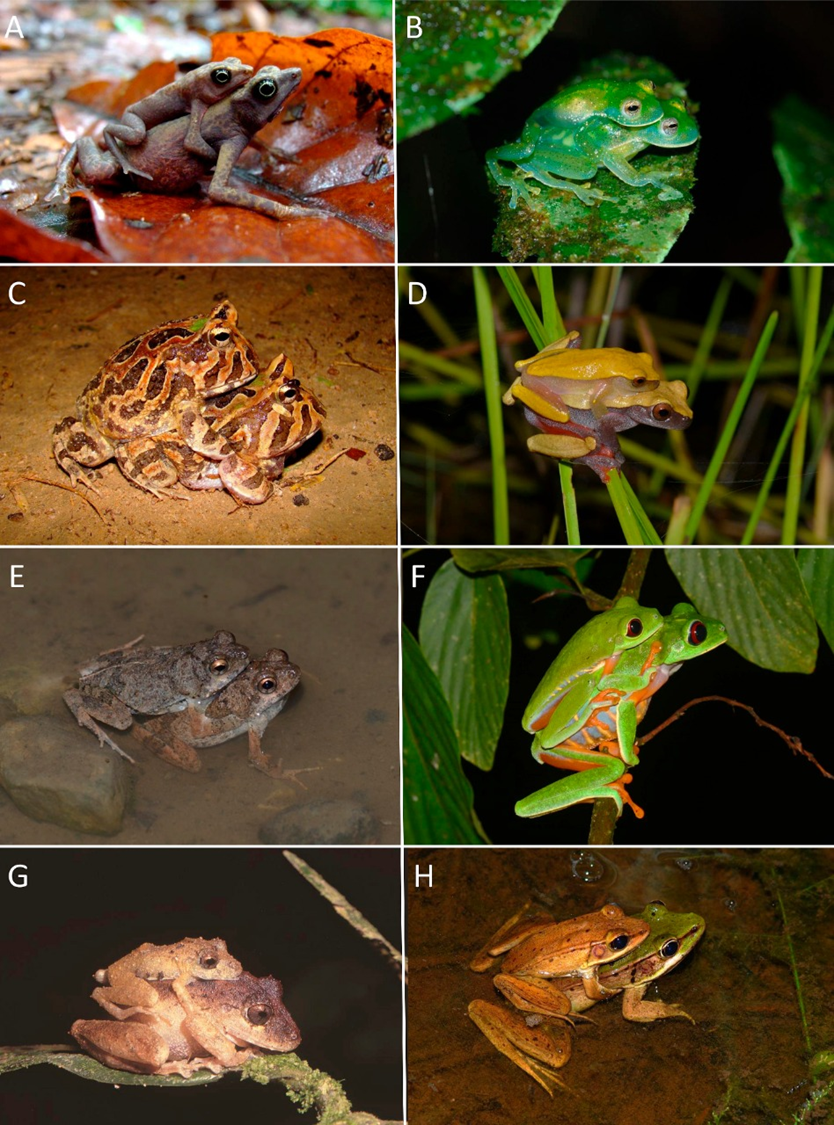
